# Temperature modifies trait-mediated infection outcomes in a *Daphnia*-fungal parasite system

**DOI:** 10.1101/2022.06.03.494706

**Authors:** Syuan-Jyun Sun, Marcin K. Dziuba, Riley N. Jaye, Meghan A. Duffy

## Abstract

One major concern related to climate change is that elevated temperatures will drive increases in parasite outbreaks. Increasing temperature is known to alter host traits and host-parasite interactions, but we know relatively little about how these are connected mechanistically – that is, about how elevated temperatures impact the relationship between epidemiologically relevant host traits and infection outcomes. Here, we used a zooplankton-fungus (*Daphnia dentifera-Metschnikowia bicuspidata*) disease system to investigate whether temperature interacted with host susceptibility traits in determining infection outcomes. We did this by exposing *D. dentifera* to *M. bicuspidata* or leaving them unexposed at either control (20°C) or warming temperatures (24°C) in a fully factorial design. We found that elevated temperatures altered the physical barrier and immune responses to parasites during the initial infection process, and that infected hosts at elevated temperatures suffered a greater reduction of fecundity and lifespan. Furthermore, the relationship between a key trait – gut epithelium thickness, which is a physical barrier – and the likelihood of terminal infection reversed at warmer temperatures. Together, our results highlight the complex ways that temperatures can modulate host-parasite interactions, and the importance of considering key host susceptibility traits when predicting disease dynamics in a warmer world.

This article is part of the theme issue ‘Infectious disease and evolution in a changing world’.

## Introduction

Elevated temperature can have major impacts on host-parasite interactions and infection outcomes, shaping disease epidemics [1]. A growing body of work demonstrates that parasites often outperform their hosts at elevated temperatures, suggesting that a warmer world will be sicker [2]. However, this outcome is not universal (e.g., [1,2]), and, at present, it is challenging to predict the impact of elevated temperature on a particular host-parasite interaction. Changing thermal regimes, such as increases in mean temperatures and extreme temperature events, can alter morphological, physiological, and chemical traits of individuals [3–5]. However, most studies have focused on the effect of climate warming on parasite development and disease transmission [5,6]; empirical tests of how temperatures mediate host traits that influence infection outcomes are still in their infancy [7].

Host traits, including physical barriers and chemical and immunological responses, play crucial roles in determining infection outcomes. Recent work suggests that decomposing different functional steps of host-parasite interactions can allow for a mechanistic understanding of how different forms of host defense mediate host susceptibility and resistance [8,9]. Susceptibility is known to vary with temperature due to changes in individual physiological and immune responses [10,11], making it so that changing temperatures can have major fitness consequences for both hosts and parasites [12,13]. Furthermore, thermal stress can induce trade-offs between parasite defenses and life history traits [14,15], such as between reproduction and infection resistance. These trade-offs can be particularly critical in sub-optimal thermal environments, such as when a host is stressed by high temperatures [14,15]. However, despite the importance of understanding how changing environments influence host susceptibility to parasites, we are still uncertain about how the traits determining host susceptibility are impacted by warmer temperatures.

The crustacean zooplankton grazers *Daphnia* sp. play a significant role in freshwater food webs, where they consume algae and are in turn consumed by larger predators [16]. *Daphnia* are also frequently infected by several microbial parasite species [17], which can impact *Daphnia* population dynamics and have cascading impacts on food webs [18]. The yeast *Metschnikowia bicuspidata* (Ascomycota: Saccharomycetales) is a generalist parasite that is commonly found infecting many *Daphnia* species (as well as other Cladocerans) in nature [17,19–22]. In stratified lakes in the Midwestern United States, infection by *M. bicuspidata* typically happens annually during late summer/early autumn, and can reach up to 60% prevalence in some populations [23,24]. The infection process begins when the parasite spores are ingested by the hosts during filter feeding. Once consumed, successful infection requires that the needle-shaped spores pierce the gut epithelium (Fig. 1), penetrating into the body cavity to develop and reproduce, and finally reaching the terminal stage when the conidia or asci fill up and kill infected hosts [9]. The next generation of spores are released from the dead hosts to the environments, where they are consumed by new hosts, completing the infection cycle [17]. In this system, host susceptibility to parasite infection is governed by two fundamental host traits: 1) a physical barrier and 2) immune defenses [9]. First, the gut epithelium functions as a barrier to spores, with epithelial thickness influencing whether spores penetrate into the body cavity [9]. Second, spores that successfully penetrate the gut epithelium (referred to as ‘hemocoel spores’ because they enter the host hemocoel after penetrating the gut) face host internal defenses, with hemocytes (immune cells) up-regulated and recruited to the site of infection ([9,25], Fig. 1b).

**Figure 1.**
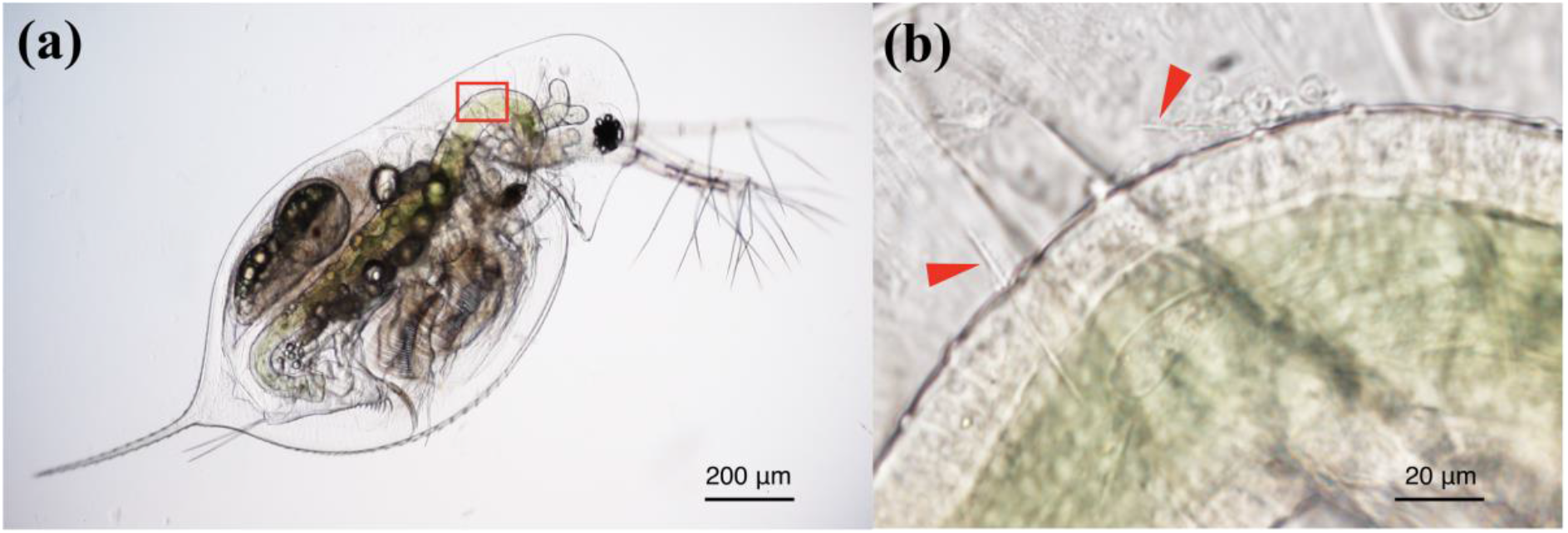
Infection process of the fungal parasite *Metschnikowia bicuspidata* within the host *Daphnia dentifera*. (a) A female host exposed to the parasite for 24 hrs. (b) *M. bicuspidata* are frequently found penetrating into the body cavity through the anterior and posterior gut epithelium. The image in (b) corresponds to the region surrounded by the red box in (a). The right and left arrows in (b) indicate hemocoel spores with and without hemocyte attachment, respectively. Photo credit: Syuan-Jyun Sun.

Here, we examine the effect of elevated temperatures on the infection process and the consequences for terminal infection outcomes. Specifically, we used the host *D. dentifera* and its fungal parasite *M. bicuspidata* as a host-parasite model system to experimentally test how elevated temperatures alter key host traits, and, in turn, how this mediates infection outcomes. *Daphnia* spp. are a well-established ecological model for climate change and disease ecology studies [16,25,26]. We began by exposing individual *D. dentifera* to *M. bicuspidata* spores in laboratory conditions at ambient temperatures (20°C) and elevated temperatures (24°C), the former being selected to match the natural epilimnetic temperature range when the epidemics start [27]. Previous work has shown that 20°C is a favorable thermal condition for optimal growth and reproduction in another species of *Daphnia* [28,29], and prior studies on *D. dentifera* have used a similar range of temperatures, since these represent ecologically relevant temperatures that are within the thermal tolerance of this species (which cannot be cultured in temperatures of 27 or higher; [7,27]). We used an average 4°C increase in temperature since it reflects a projected climate change scenario by the end of this century [30], with summer lake surface temperatures having risen at a rate of 0.34°C per decade since 1985 [31]. During the infection stage, we investigated how temperature modified key host susceptibility traits including gut epithelium thickness and hemocyte responses to attacking spores. We also evaluated the fitness consequences of temperature and terminal infection (defined as the presence of asci in the host hemolymph; [9]) on both host and parasite. We hypothesized that host susceptibility traits, including gut epithelium thickness and immune responses, determine the infection process, and that temperature will influence these traits during infection, shaping the likelihood of terminal infection. More specifically, we predicted that elevated temperatures would hinder physical barrier and/or immune defense against parasites, thus resulting in increased terminal infection and reduced host fitness.

## Methods

### (a) Study species: origin and maintenance

*Daphnia dentifera* is a cyclically parthenogenetic crustacean commonly found in freshwater lakes in temperate North America [24]. *Metschnikowia bicuspidata* is an ascomycete yeast that commonly causes epidemics in *D. dentifera* [24]. We used the “Standard” genotype of *D. dentifera* (originally isolated from a lake in Barry County, Michigan) and the “Standard” isolate of the fungal parasite *M. bicuspidata* (Baker Lake in Barry County, Michigan). The *D. dentifera* stock was maintained as small populations in 150 mL beakers (5 animals per beaker, 40 beakers in total) filled with 100 mL filtered lake water. All animals were fed three times a week with phytoplankton food (*Ankistrodesmus falcatus*, 20,000 cells/mL). Prior to the experiments, *Daphnia* were reared under standardized lab conditions (on a 16:8 photoperiod at 22°C) for three generations. The *M. bicuspidata* culture was maintained by infecting 75 *D. dentifera* (two weeks to a month old) with *M. bicuspidata* (using spores obtained by grinding up infected *D. dentifera* that had been collected 1-2 weeks prior and stored in a refrigerator before use) at a density of 250 spores/mL in a 1000 mL Erlenmeyer flask filled with 900 mL filtered lake water. The culture was fed three times a week (*A. falcatus*, 20,000 cells/mL) and checked every 10 days for infected *D. dentifera*, which were then placed in a 1.5 mL tube of 100 μL filtered lake water and stored in a refrigerator before use.

### (b) Experimental design

To test the effect of increased temperatures on our host-parasite system, we used two temperatures (20°C vs. 24°C) and the presence/absence of the parasite in a fully factorial design with 55 and 45 replicates in the presence and absence of the parasite, respectively, per temperature treatment. We used a larger number of animals in parasite present treatment to accommodate potential mortality rate associated with spore checking at initial infection stage (see below). We collected neonates aged 1-2 days old from our laboratory stock and reared them at either 20°C or 24°C, maintaining each temperature treatment in separate incubator (I-41VL, Percival Scientific). Each juvenile was kept individually in a 50 mL beaker filled with 50 mL filtered lake water and fed three times a week (20,000 cells/mL *A. falcatus*). For the parasite-exposed treatment, we exposed juveniles that were 5-6 days old to parasite spores; control treatment animals were left unexposed. To control for variation in developmental stage due to temperature differences during parasite exposure (i.e., faster growth at 24°C), we used a degree-day approach (accumulated product of time and temperature [32,33]) to expose *Daphnia* to parasites at an age of 5 and 6 days, for 24°C and 20°C, respectively. This allowed us to minimize potential body size differences between temperature treatments (Χ^2^ = 2.69, d.f. = 1, *P =* 0.101) during parasite exposure, since larger body size leads to higher spore intake [34]. Therefore, at degree-day 120 (i.e., 5 days x 24 °C or 6 days x 20°C), we inoculated experimental animals with a suspension of spores obtained by crushing 45 *D. dentifera* infected with *M. bicuspidata*. Using a hemocytometer, we determined the spore density used (145 spores/mL), which is biologically relevant to field populations [9]. Meanwhile, the unexposed animals received a placebo solution containing the same amount of tissue from uninfected *D. dentifera*. Animals were exposed to parasites or placebo for 24 hr; these animals were fed 20,000 cells/mL *A. falcatus* and kept at 16:8 light:dark cycle. Experimental animals were then transferred individually to beakers filled with 50 mL filtered lake water, fed three times a week (20,000 cells/mL *A. falcatus*) and maintained at 16:8 light:dark cycle until the end of the experiment. The experiment was conducted in two blocks between January and March 2022.

### (c) Data collection

To determine the impact of temperature on the earliest stages of the infection process, we examined the hosts at the end of the 24 hour inoculation period under an Olympus BX53F compound microscope (200-400X magnification). We scanned for the presence of spores in the anterior and posterior regions of the host gut and body cavity, where spores are most likely to penetrate [9,35] (see also Fig. 1). Ingested spores were categorized into the two classes (*sensu* [9]): embedded spores (i.e., spores partially embedded in the gut epithelium) and hemocoel spores (i.e., spores that had successfully penetrated into the body cavity). We were interested in quantifying the extent to which the gut is a barrier to infecting spores; we determined ‘gut resistance’ as the proportion of attacking spores (embedded spores + hemocoel spores) that were blocked by the gut barrier (i.e., embedded spores divided by attacking spores). To quantify the immune response, we counted the total number of hemocytes attached to hemocoel spores (Fig. 1b). We also determined the number of hemocytes per spore by dividing the total number of hemocytes by the number of hemocoel spores.

As we quantified the immune responses in infected individuals, we also imaged the gut epithelium at high magnification (400X) and measured the height of anterior midgut epithelial cells at the 90-degree bends in the C-shaped gut (Fig. 1), from where we determined the average gut epithelium thickness by haphazardly selecting three epithelial cells; the gut epithelium in *Daphnia* is one cell layer thick. We also measured body size by drawing a straight line from the center of the eye to the base of the spine. All measurements were made using the cellSens Standard Software (Olympus, version 1.18). In total, 50 and 53 individuals for control and warming temperatures were successfully measured.

To determine fitness components of the host, we checked all individuals daily for mortality and counted the number of offspring produced, which were then removed from the beakers. We estimated fecundity (total number of offspring) for all animals 20 days after exposure, when the last infected host was found dead. Upon death, each infected host was placed in a 1.5 mL tube of 100 μL deionized water and stored in a refrigerator for further spore yield counts. To determine spore yield per host, we ground the host using an electric pestle for 60 seconds, and mixed the spore solution well before adding a 10 μL sample to a Neubauer Hemocytometer. We estimated the spore yield by averaging the number of mature spores from four grids. We excluded animals that died within 7 days after exposure due to early mortality, during which infection status cannot be accurately diagnosed, resulting in a total of 162 individuals (control temperatures: *n* = 37 and 38 for control and parasite treatment; warming temperatures: *n* = 44 and 43 for control and parasite treatment).

### (d) Statistical analyses

We analyzed the data using generalized linear mixed models (GLMM) with the glmer function in the *lme4* package [36] in R version 4.1.2 [37]. Analysis of variance (ANOVA) was performed in the *car* package [38], with type III sums-of-squares. Model selection was conducted with a stepwise regression approach based on the Akaike information criterion by removing non-significant interactions. Once a significant interaction term was detected, Tukey *post-hoc* comparisons were made to assess differences among individual treatments in the *emmeans* package [39]. For all of the models described below, block was included as a random effect.

We analyzed the effects of temperature on gut epithelium thickness with a Gaussian distribution by including temperature treatment and body size as fixed effects. To investigate if guts were a barrier to spore infection, we analyzed gut resistance with a Gaussian distribution by including two fixed effects: temperature treatment and gut epithelium thickness, which was better fitted as a second-degree polynomial term. We analyzed the number of hemocoel spores in a similar manner, including temperature treatment and gut epithelium thickness as fixed effects, but with a Poisson distribution.

We analyzed the effect of temperature on the total number of hemocytes with a negative binomial GLMM to account for data overdispersion. We included temperature treatment and the number of hemocoel spores as fixed effects. We further analyzed the number of hemocytes per spore with a Gaussian distribution by including temperature treatment and gut epithelium thickness as fixed effects. Because a significant interaction was detected, we also analyzed each temperature treatment separately to evaluate how the number of hemocytes per spore varied with gut epithelium thickness.

We investigated how temperature treatments and host susceptibility traits jointly affected terminal infection outcomes. We first quantified their impact on parasite infectivity (terminal infection: 1; no terminal infection: 0), where probability of successful infection was analyzed with a binomial distribution and logit link function. We also analyzed effects of temperature and host traits on spore yield per host; this data was log transformed prior to analysis with a Gaussian distribution to meet the assumption of normality for regressions. In both analyses, we included temperature treatment, gut epithelium thickness, and number of hemocytes per spore as fixed effects. Because of significant interactions between temperature treatment and gut epithelium thickness, we further analyzed each temperature treatment separately to evaluate differential effects of gut epithelium thickness on parasite infectivity and spore yield per host.

We investigated the impact of temperature and infection on host fitness, in terms of survival and fecundity. Host survival was analyzed with a Cox proportional hazard mixed effect model in the *coxme* package [40]. We included temperature (control/warming) and parasite treatment (control/exposed/infected) as fixed effects and included block as a random effect. We used three categories related to parasite treatment to evaluate whether hosts that were exposed to parasites but that did not develop terminal infections (referred to as “exposed” in our analyses) differed from uninfected controls and/or from hosts that were exposed and developed terminal infections (referred to as “infected” in our analyses). The data was censored to indicate *Daphnia* that remained alive at the end of the study. Fecundity was analyzed with a negative binomial GLMM to account for data overdispersion, with temperature and parasite treatments included as fixed effects.

## Results

*Daphnia* that experienced higher temperatures had gut epithelial cells that were, on average, 16% thicker than those at ambient temperatures (control: 18.78 ± 0.46 μm, warming: 21.76 ± 0.37 μm; Fig. 2a), even when controlling for body size (Fig. 2b). Both thinner and thicker gut epithelia acted as barrier resistance to attacking spores – that is, a smaller proportion of attacking spores were successfully blocked in individuals with intermediate gut epithelium thickness (Table 1; Fig. 2c), leading to a larger number of hemocoel spores (Table 1; Fig. 2d); there was no difference in gut resistance or hemocoel spores between the temperature treatments (Table 1).

**Table 1.** Results from the final models analyzing the traits associated with infection processes.

| **Dependent variable** | **Explanatory variables** | **X<sup>2</sup>** | **d.f.** | **p value** |
| --- | --- | --- | --- | --- |
| Gut resistance | Gut thickness | 0.47 | 1 | 0.491 |
|  | Gut thickness<sup>2</sup> | 6.21 | 1 | **0.013** |
|  | Temperature | 0.00 | 1 | 0.971 |
| Hemocoel spore | Gut thickness | 1.79 | 1 | 0.181 |
|  | Gut thickness<sup>2</sup> | 5.25 | 1 | **0.022** |
|  | Temperature | 0.08 | 1 | 0.771 |
| Total recruited hemocytes | Hemocoel spore | 44.61 | 1 | **<0.001** |
|  | Temperature | 16.40 | 1 | **<0.001** |
| Hemocytes per spore | Gut thickness | 4.94 | 1 | **0.026** |
|  | Gut thickness<sup>2</sup> | 8.51 | 1 | **0.004** |
|  | Temperature | 5.28 | 1 | **0.022** |
|  | Gut thickness x Temperature | 2.92 | 1 | 0.087 |
|  | Gut thickness<sup>2</sup> x Temperature | 6.10 | 1 | **0.013** |

Statistically significant *p* values are highlighted in bold.

**Figure 2.**
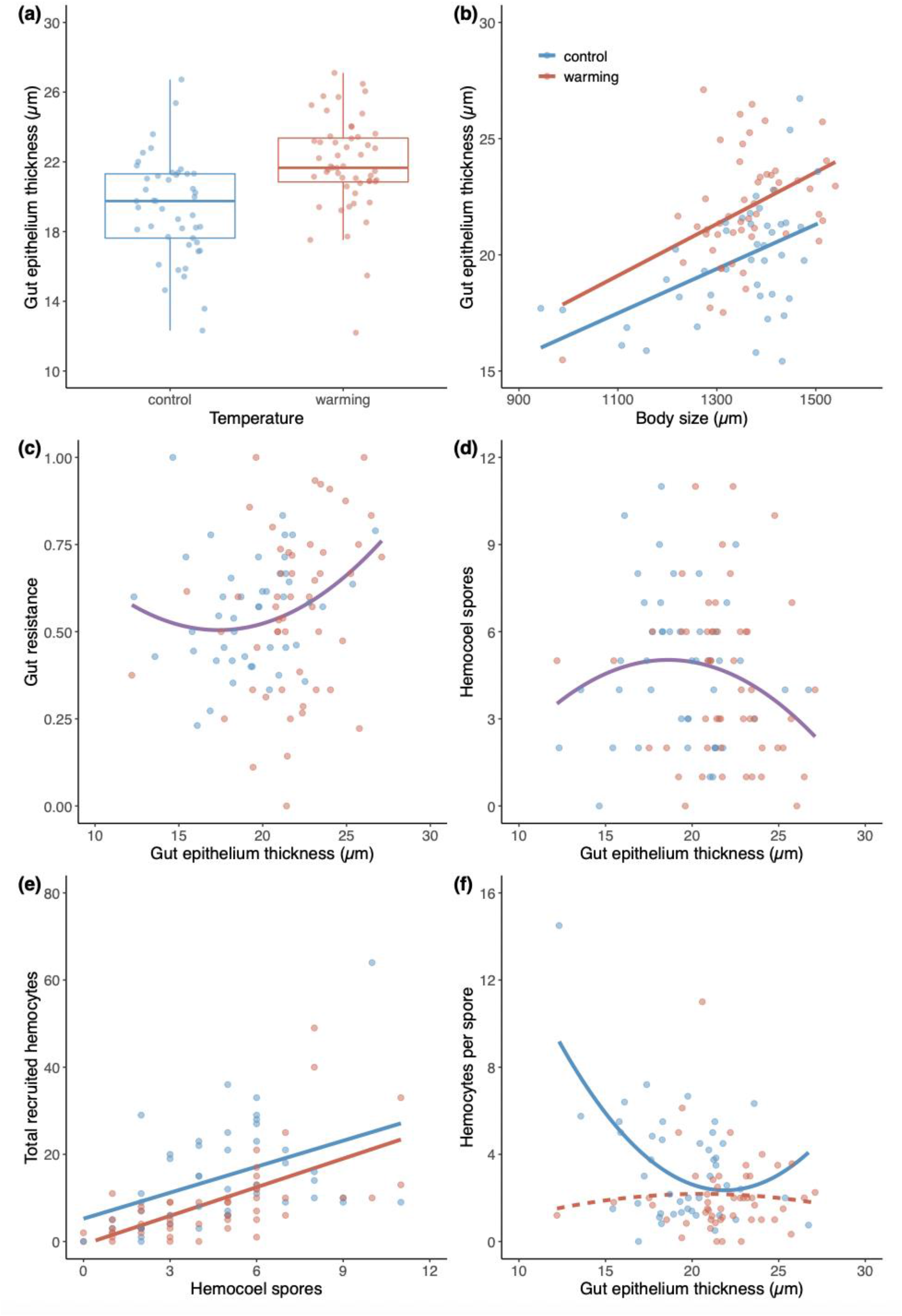
The effects of temperatures on two different forms of defense (i.e., barrier and immunity) and their relationship with each other. (a) The gut epithelium, which serves as a barrier to parasite entry, is thicker in animals that experienced warmer temperatures, (b) even in animals of the same body size. (c) Animals with moderately thick guts had the lowest gut resistance, regardless of temperature, leading to (d) the number of hemocoel spores being highest for intermediate gut epithelium thickness; ‘gut resistance’ was quantified by dividing the number of embedded spores by the total number of attacking spores (that is, by the number of embedded spores + hemocoel spores), so the inverse relationship between gut resistance and hemocoel spores is expected. (e) Hemocoel spore number positively predicted total number of recruited hemocytes, with control temperatures consistently having more hemocytes than warming temperatures. (f) The relationship between gut epithelium thickness and immune responses, measured as the number of hemocytes per hemocoel spore, differed between the two temperatures. The box plot in (a) shows median values, the 25^th^ and 75^th^ percentiles, and interquartile ranges. Solid and dashed lines indicate statistically significant and non-significant regressions, respectively, predicted from GLMMs. Each panel also plots the raw data points (each from a single animal).

Turning to immune responses, we found that both the number of hemocoel spores and the temperature impacted the number of hemocytes that attacked spores (Table 1, Fig. 2e): as more spores penetrated into the body cavity, the total number of hemocytes that attached to the infecting spores increased, and there were generally more hemocytes in ambient temperatures compared to elevated temperatures. Consequently, warming resulted in consistently lowered number of hemocytes per spore that successfully penetrated, compared to ambient temperatures (Table 1; Fig. 2e). We also found that number of hemocytes per spore was explained by gut thickness, but the relationship differed between temperature treatments (Table 1; Fig. 2f). Specifically, thinner gut epithelia were associated with an increasing number of hemocytes per spore under ambient temperatures (Χ^2^ = 10.20, d.f. = 1, *P =* 0.001); under warming, this relationship was dampened, with generally fewer hemocytes per spore and no significant relationship between gut epithelium thickness and hemocytes per spore (Χ^2^ = 0.01, d.f. = 1, *P =* 0.905).

We found that probability of terminal infection was predicted by gut thickness, but the patterns differed markedly between temperature treatments (Table 2; Fig. 3a). Specifically, the probability of terminal infection increased as gut epithelium thickness increased at ambient temperatures (Χ^2^ = 4.74, d.f. = 1, *P =* 0.029), whereas the probability of terminal infection decreased as gut epithelium thickness increased at higher temperatures (Χ^2^ = 4.08, d.f. = 1, *P =* 0.043). While *Daphnia* with thicker guts had similarly high probability of infection irrespective of temperature treatment (Χ^2^ = 0.70, d.f. = 1, *P =* 0.401), those with thinner guts tended to have relatively lower probability of infection at control than warming temperatures (Χ^2^ = 5.50, d.f. = 1, *P =* 0.019). For *Daphnia* that developed terminal infection, gut thickness (Χ^2^ = 0.08, d.f. = 1, *P =* 0.779) had no effect on spore production, and there was no difference between temperature treatments (Χ^2^ = 1.85, d.f. = 1, *P =* 0.173). As a result, the overall effect of gut thickness on spore production of animals that had been exposed to infection was mediated by temperature treatments (Table 2; Fig. 3b), due to their effect on the likelihood of terminal infection; of all animals that were exposed, more spores were produced in hosts with thicker guts only at ambient temperatures (Χ^2^ = 5.93, d.f. = 1, *P =* 0.015) but not at warming temperatures (Χ^2^ = 5.19, d.f. = 1, *P =* 0.023; Fig. 3c). *Daphnia* with thicker guts had similarly high number of spores produced in both control and warming temperatures (Χ^2^ = 1.04, d.f. = 1, *P =* 0.309), whereas *Daphnia* with thinner guts produced fewer spores at control than at warming temperatures (Χ^2^ = 5.99, d.f. = 1, *P =* 0.014; Fig. 3d).

**Table 2.** Results from the final models analyzing the parasite fitness components of terminal infection outcomes.

| **Dependent variable** | **Explanatory variables** | **$\chi^2$** | **d.f.** | ***p* value** |
| --- | --- | --- | --- | --- |
| Probability of terminal infection | Gut thickness | 2.68 | 1 | 0.102 |
|  | Temperature | 7.17 | 1 | **0.007** |
|  | Hemocytes per spore | 3.06 | 1 | 0.080 |
|  | Gut thickness x Temperature | 6.53 | 1 | **0.011** |
| Spore yield per host | Gut thickness | 2.20 | 1 | 0.138 |
|  | Temperature | 6.77 | 1 | **0.009** |
|  | Hemocytes per spore | 3.12 | 1 | 0.077 |
|  | Gut thickness x Temperature | 5.77 | 1 | **0.016** |

Statistically significant *p* values are highlighted in bold.

**Figure 3.**
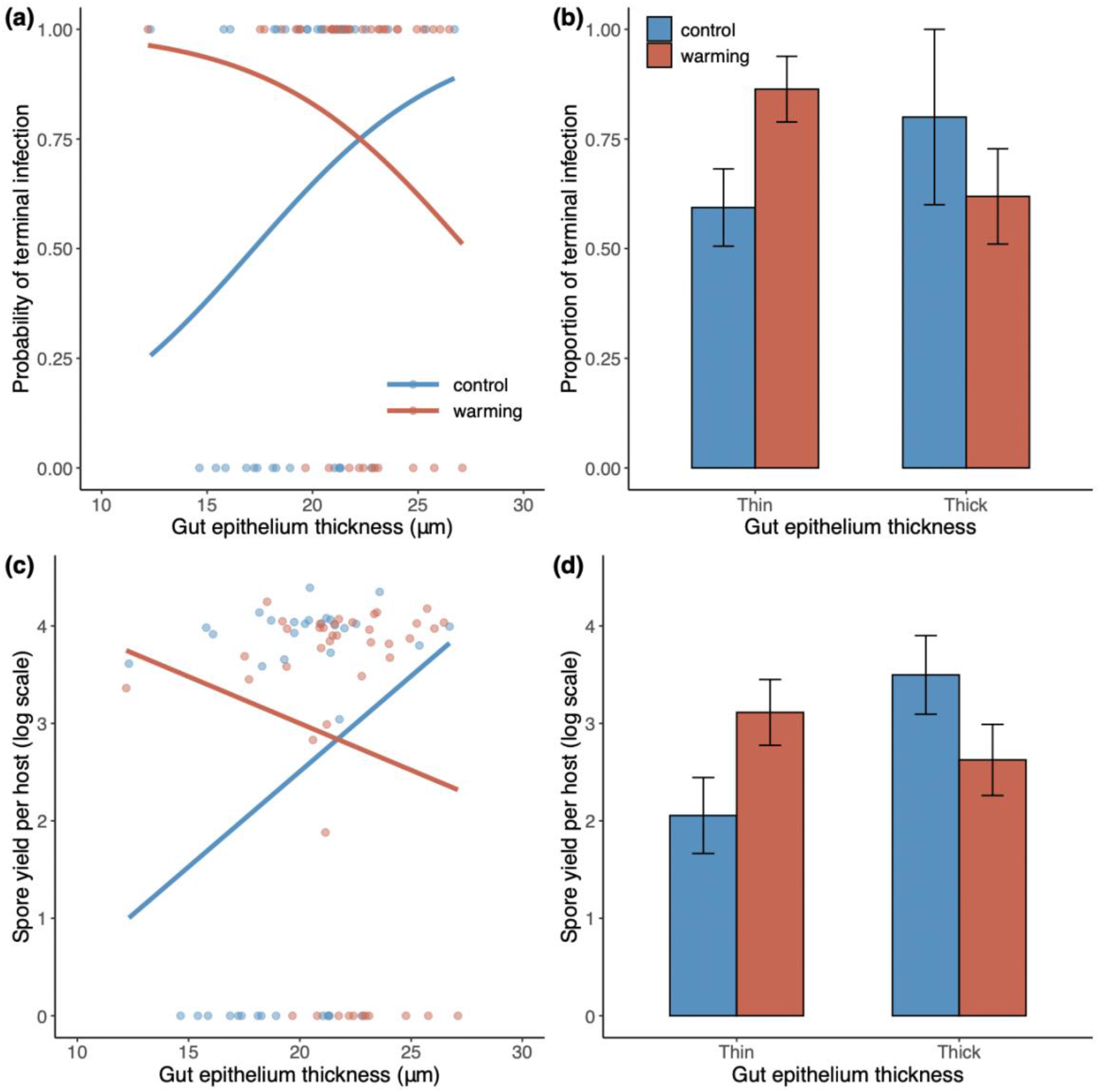
(a,b) The probability of terminal infection and (c,d) spore yield per host in relation to gut epithelium thickness at control and warming temperatures. Significant interactions between temperature treatments and gut epithelium thickness were detected, and so the data were split into two parts in (b,d), based on the intersection values of gut epithelium thickness in (a,c) to evaluate the effects of temperature. The lines indicate statistically significant regressions predicted from GLMMs. Means and standard error bars are shown.

*Daphnia* fitness was impacted by both parasite treatment and temperature (Fig. 4). *Daphnia* exposed to *Metschnikowia* suffered from reduced fecundity but the extent depended upon temperature treatments (Temperature x Parasite treatments: Χ^2^ = 20.76, d.f. = 2, *P <* 0.001; Fig. 4a). Specifically, *Daphnia* successfully infected by *Metschnikowia* experienced fecundity reduction compared to unexposed *Daphnia* at control temperatures (compare the blue “control” bar and the blue “infected” bar in Fig. 4a; *post-hoc* comparison control v. infected: *z* = 8.11, *P <* 0.001), but there was greater reduction at warming temperatures (red “control” bar *vs*. red “infected” bar in Fig. 4a; *post-hoc* comparison control v. infected: *z* = 15.83, *P <* 0.001), resulting in lower reproductive success of infected *Daphnia* at warming than at control temperatures (blue “infected” bar *vs*. red “infected” bar in Fig. 4a; *post-hoc* comparison control v. warming: *z* = 3.12, *P =* 0.002). For both temperature treatments, *Daphnia* that were exposed but uninfected had intermediate numbers of offspring – they produced more offspring than infected animals but fewer than the control (“exposed” treatment in Fig. 4a); no difference was found between the two temperature treatments (*post-hoc* comparison control v. warming: *z* = -0.15, *P =* 0.878). Host survival was lower at warm than at control temperatures, and survival was highest in unexposed controls, followed by exposed but uninfected, and lowest in infected animals (Fig. 4b). Host survival was independently affected by temperature (Χ^2^ = 28.40, d.f. = 1, *P <* 0.001) and parasite treatments (Χ^2^ = 85.93, d.f. = 2, *P <* 0.001), with the lowest overall survival for animals that were infected at warm temperatures.

**Figure 4.**
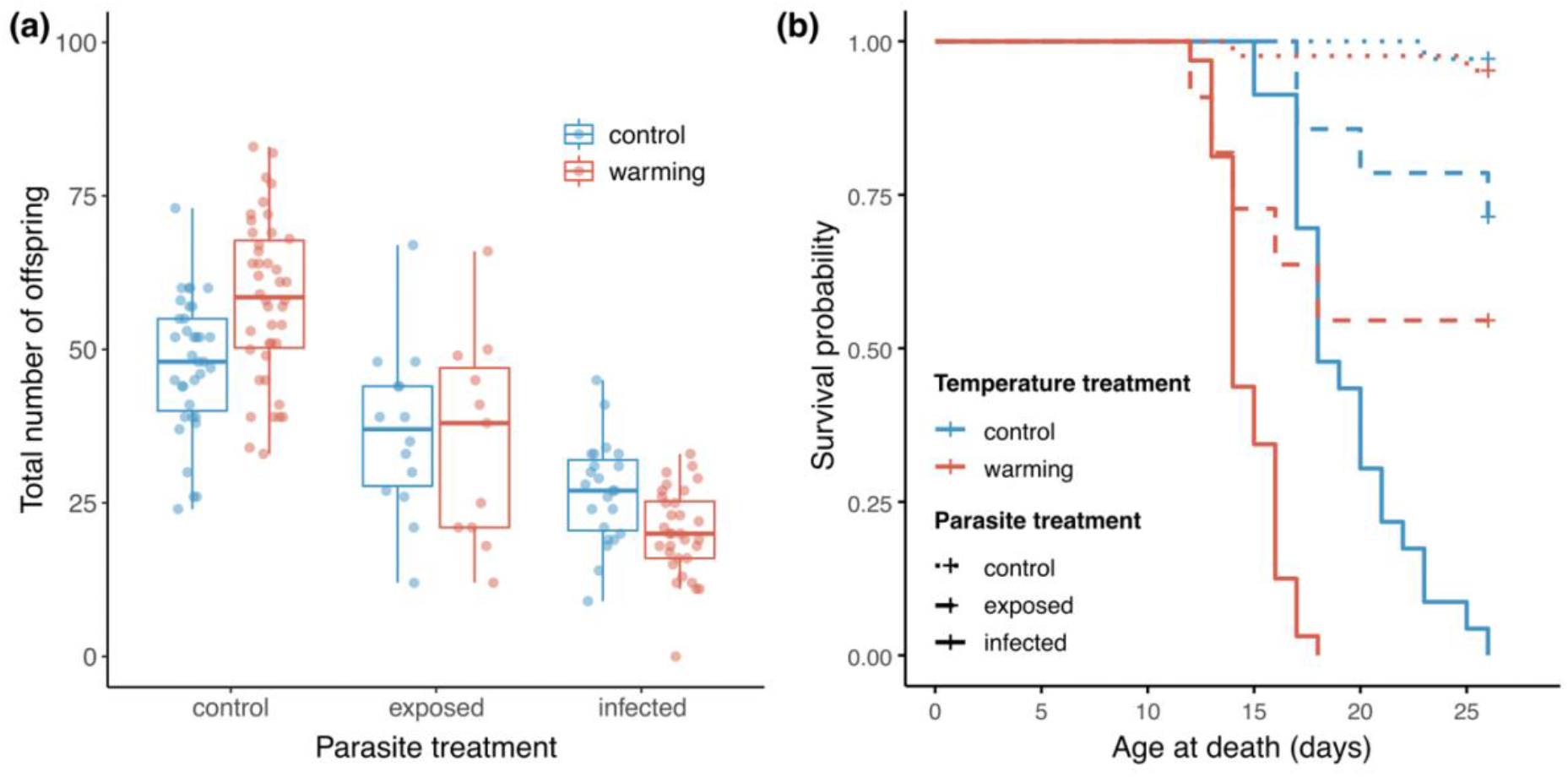
The effect of temperature change and *Metschnikowia* infection on *Daphnia* fitness components, measured as (a) fecundity (i.e., total number of offspring) and (b) survival, i.e., the proportion of individuals remaining alive at each time. In parasite treatment, ‘control’ represents hosts unexposed to parasites, whereas ‘exposed’ and ‘infected’ represents hosts exposed to parasites but that remained uninfected or became infected, respectively. The box plots show median values, the 25^th^ and 75^th^ percentiles, and interquartile ranges. Solid lines represent the infected individuals, dashed lines represent exposed, and dotted lines represent uninfected individuals.

## Discussion

Our results showed that elevated temperatures altered physical barriers and immune responses to infection and exacerbated the effects of the fungal parasite *Metschnikowia bicuspidata* on the host *Daphnia dentifera* – while hosts at elevated temperatures had (beneficial) thicker guts, they also had dampened immune responses, and infected hosts suffered a greater reduction in fecundity and lifespan at elevated temperatures. Moreover, we found that the relationship between a key physical barrier, gut epithelium thickness, and the probability of developing a terminal infection was qualitatively different at elevated temperatures: at ambient temperatures, individuals with thicker guts were more likely to develop terminal infections, but at warmer temperatures, individuals with thinner guts were more likely to develop terminal infections. Our findings demonstrate that we must understand how temperature influences key traits and their relationship with infection outcomes to predict the ecological and evolutionary consequences of host-parasite interactions in a warmer world.

In systems where immune responses are largely modulated by environmental temperature (e.g., ectotherms), elevated temperature can interact with immune functions in complex and unpredictable ways [41]. Temperature increase is known to cause immune suppression in both terrestrial and aquatic systems [42], and developing a mechanistic understanding of how temperature modulates host immune responses and host-parasite interactions has become a central focus in ecological immunity [41,43,44]. We found that temperature modulates host-parasite interactions through associated changes in key host susceptibility traits, which were differentially affected by temperatures. Previous work in this system found that the gut epithelium acted as a barrier to *M. bicuspidata* infection, with thinner guts being more likely to prevent spore penetration [9]. Our findings corroborate this within the same range of gut epithelium thickness (10-20 *μ*m). However, we found an overall u-shaped relationship between gut epithelium thickness; beyond ∼20 *μ*m, thicker guts were more resistant to attacking spores (Fig. 2b). Because of this relationship between gut thickness and resistance to attacking spores, we found fewer hemocoel spores at both thinner and thicker ends of the gut epithelium gradient (Fig. 2c). This shows that it is not sufficient to just measure how key traits might change in an altered environment; if we had just measured gut thickness, we would have concluded, because individuals raised at warmer temperatures had thicker guts, that they would also be more susceptible to infection. Instead, because the relationship between that key trait and infection also changed in warmer environments, the observed increase in gut thickness under elevated temperatures helped hosts resist attacking parasites at early infection stages.

At elevated temperatures, spores that successfully penetrated into the host’s body cavity encountered reduced immune responses: for a given number of hemocoel spores, hosts generally recruited fewer hemocytes at warmer temperatures than the control (Fig. 2e). At control temperatures, hosts with the thinnest gut epithelia recruited the most hemocytes per spore, suggesting a potential trade-off between immune functions and physical barriers. A trade-off between different types of immune functions and life-history traits has also been widely found in other systems [45–47]. However, this relationship between gut thickness and hemocyte response was not present at warmer temperatures, possibly because there were too few hemocytes per spore for any relationship to be apparent. This reduced immune response might indicate that immunity is costly to maintain at warmer temperatures, and/or might have not occurred because energy was diverted to other life history traits, such as thicker gut barriers. This temperature-mediated trade-off in immunity is supported by a study on the reef coral *Porites cylindrica*, which was challenged with injury at ambient and warmer temperatures [48]; warming induced *Porites cylindrica* to divert resources away from immune response to ensure homeostasis [48]. Hemocytes are essential components of immune responses in invertebrates, and in some cases, can prevent infection development [42]. Nevertheless, elevated hemocyte number in our system did not necessarily mean *Daphnia* were more resistant to the parasites. Instead, accumulating evidence suggests that increasing hemocyte number can be a symptom of infection, rather than an effective immune response [9,49]. Our terminal infection results provide some support for this: hosts with more hemocytes per spore did not have lower terminal infection likelihood or spore yield; instead, there was either no significant relationship or a slightly positive one between hemocytes per spore and terminal infection and spore yield (Table 2; Fig. S1).

Preventing parasite infection is pivotal because infected hosts suffered from shortened lifespan and fecundity loss, particularly under warming (Fig. 4). We found that 39.5% of the exposed hosts at control and 25.6% of the exposed hosts at warming temperatures did not develop terminal infection, and that they outperformed those who developed terminal infections (Fig. 4). Nevertheless, at both control and warmed temperatures, *Daphnia* that were exposed to the parasite but did not develop terminal infections still performed worse than those who had not encountered parasites, suggesting that fighting infection is costly; most, but not all, of the “exposed” animals in our study were observed to have hemocoel spores 24 hours after the end of the exposure period and, therefore, successfully fought off early infections. The probability of terminal infection was higher under warming, but interestingly the outcomes depended upon gut epithelium thickness. Indeed, we found contrasting patterns of the effect of gut epithelium thickness on infection outcomes, which were context-dependent upon temperatures. At control temperatures, thicker gut epithelia predicted higher infectivity, but at warming temperatures thicker gut epithelia instead inhibited parasite fitness (Fig. 4). Consequently, it was the *Daphnia* with thinner gut epithelia that were more susceptible to parasite infection under warming. Although the gut epithelium has been widely found to function as a physical barrier to ingested parasites [9,42,50,51], how environmental factors influence gut resistance to parasite infection remains unexplored. This finding, to our knowledge, provides the first evidence of a temperature-driven shift in physical barriers against parasites. Further investigation is needed to identify the exact mechanisms underlying the regulation of key host susceptibility traits in response to temperature changes.

In conclusion, our results demonstrate that temperature modulates the infection process and the outcome of host-parasite interactions in complex ways. By tracking host susceptibility traits from initial stages of the infection process through terminal infection outcomes, our results show that temperature affects not only key host susceptibility traits and the outcomes of infection, but also the relationship between those traits and infection outcomes. Thus, our study highlights the importance of examining key functional traits associated with the infection process, and understanding how relationships between those traits and infection change with increased temperatures, to better understand host-parasite interactions and disease dynamics in a changing world.

## Acknowledgement

We thank members of the Duffy Lab for logistic support, particularly Kira Monell and Siobhan Calhoun for the maintenance of *Daphnia* and *Metschnikowia* cultures. This work was supported by the Gordon and Betty Moore Foundation (GBMF9202; DOI: https://doi.org/10.37807/GBMF9202).

## Supplemental Material

**Figure S1.**
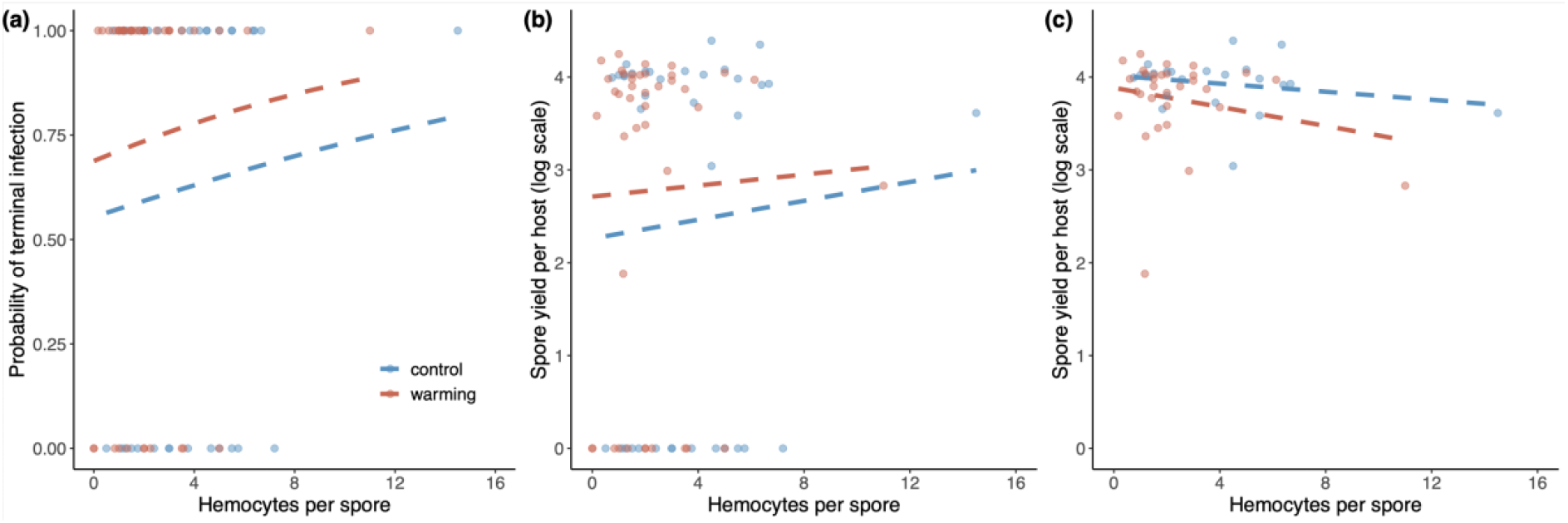
The probability of (a) terminal infection and (b) spore yield per host in relation to the number of hemocytes per spore at control and warming temperatures. (C) Considering just *Daphnia* that developed terminal infection, hemocyte number per spore had no effect on spore production (Χ^2^ = 0.11, d.f. = 1, *P =* 0.739); points shown in (c) are a subset of the points shown in (b). Once the two outliers were removed (hemocytes per spore > 8), the effects of hemocytes per spore remained non-significant when predicting the probability of terminal infection (Χ^2^ = 0.82, d.f. = 1, *P =* 0.365), spore yield per host (Χ^2^ = 1.32, d.f. = 1, *P =* 0.251), and spore yield per host that developed terminal infection (Χ^2^ = 1.10, d.f. = 1, *P =* 0.296). The lines indicate statistically non-significant regressions predicted from GLMMs.

